# Characterising indel diversity in a large *Mycobacterium tuberculosis* outbreak – implications for transmission reconstruction

**DOI:** 10.1101/2022.10.26.513840

**Authors:** Benjamin Sobkowiak, Caroline Colijn

**Affiliations:** Department of Mathematics, Simon Fraser University, Burnaby, Canada

**Keywords:** whole genome sequencing, *Mycobacterium tuberculosis*, pathogen transmission, phylogenetics, molecular evolution

## Abstract

Genomic sequencing of *Mycobacterium tuberculosis (Mtb)*, the primary aetiological agent of tuberculosis (TB) in humans, has been used to understand transmission dynamics and reconstruct past outbreaks. Putative transmission events between hosts can be predicted by linking cases with low genomic variation between pathogen strains, though typically only variation in single nucleotide polymorphisms (SNPs) is used to calculate divergence. In highly clonal *Mtb* populations there can be many strains that appear identical by SNPs, reducing the utility of genomic data to disentangle potential transmission routes in these settings. Small insertions and deletions (indels) are found in high numbers across the *Mtb* genome and can be an important source of variation to increase the observed diversity in outbreaks. Here, we examine the value of including indels in the transmission reconstruction of a large *Mtb* outbreak in London, UK, characterised by low levels of SNP diversity between 1998 and 2013. Our results show that including indel polymorphism decreases the number of strains in the outbreak with at least one other identical sequence by 43% compared to using only SNP variation and reduces the size of largest clonal cluster by 53%. Considering both SNPs and indel polymorphisms alters the reconstructed transmission network and decreases likelihood of direct transmission between hosts with variation in indels. This work demonstrates the importance of incorporating indels into *Mtb* transmission reconstruction and we provide recommendations for further work to optimise the inclusion of indel diversity in such analyses.

## Introduction

Tuberculosis (TB), caused by the bacteria *Mycobacterium tuberculosis (Mtb)*, remains a global health concern, with an estimated 1.5 million TB deaths in 2020 (World Health Organization 2021). Recent positive trends in reducing global TB incidence as part of the World Health Organization (WHO) ‘End-TB’ strategy (World Health Organization 2017) have been set back by the COVID-19 pandemic, resulting in fewer newly diagnosed TB cases and a higher number of TB deaths compared to 2019 (World Health Organization 2021). These worsening trends are projected to continue, and work is needed to better understand the biological and epidemiological drivers of *Mtb* transmission to achieve targets to eliminate TB globally. Advances in next-generation sequencing technologies and the availability of whole genome sequencing (WGS) has allowed genomic data to become a routine tool for characterising evolutionary relationships between strains. Genomic divergence can be used in combination with epidemiological information to reconstruct putative transmission networks (Didelot et al. 2017; Klinkenberg et al. 2017) and give insights into the transmission dynamics within an outbreak, including who infected whom and the timing of transmission events (Gardy et al. 2011; Romanowski et al. 2020; Sobkowiak et al. 2020; Xu et al. 2020).

Typically, divergence is measured using single nucleotide polymorphisms (SNPs), *de novo* point substitutions, which are the primary source of variation in *Mtb* genomes. Phylogenetic trees built using variation in SNPs have been used to describe evolutionary relationships and reconstruct transmission histories using *Mtb* WGS data ((Hatherell et al. 2016; Ayabina et al. 2018; Sobkowiak et al. 2020)), though relatively low mutation rates in TB often means that there is limited SNP diversity within outbreaks, especially over short timescales (Lee et al. 2020). This can complicate transmission reconstruction particularly when two or more strains appear genetically identical, requiring validation with epidemiological tracing to resolve the correct direction and source of transmission; the necessary data may not be available. Indels (short insertions and deletion of up to 50 bases) are an important source of *Mtb* genomic variation that are often excluded from comparative analysis due to issues assembling these regions from short-read sequence data. Indels are mainly found in intergenic regions, though indel polymorphisms that modify gene functions have been identified, notably in the PE/PPE gene families and loci associated with antimicrobial resistance and adaptive plasticity (Coll et al. 2014; Phelan et al. 2019; Gupta and Alland 2021). The frequency of indels within *Mtb* genomes suggests that they can be a valuable source of information for detecting genomic diversity between closely related strains.

Understanding the evolution and diversity of indels will should allow us to increase the observable variation in *Mtb* outbreaks and resolve evolutionary relationships between strains with limited SNP diversity. In this study, we examined the evolutionary and temporal signal in indels in WGS isolates, including serially sampled hosts, from large *Mtb* outbreak in London, UK between 1998 – 2013 (Xu et al. 2020), a persistent outbreak characterised by low levels of genetic diversity and pan-resistance to isoniazid (INH), with low levels of resistance to other antimicrobials. We investigated the SNP and indel diversity of the outbreak strains and compared the transmission events inferred with a SNP-only analysis with an analysis including indels using TransPhylo (Didelot et al. 2017), a Bayesian approach for reconstructing transmission networks from phylogenetic trees. To our knowledge, this is the first comprehensive study to use variation in indels to probabilistically reconstruct transmission in a complex TB outbreak. We examine the potential utility of including indels as genetic markers in *Mtb* genomic analysis in this context.

## Results

### Indel polymorphism and temporal signal

Sequence analysis of 332 *Mtb* isolates collected the London, UK outbreak revealed 1518 SNP and 197 indel variants called against the H37Rv reference strain. The majority of these indel polymorphisms were found in coding regions (152/197; 77%), with a high number found in PE/PPE genes (63/197; 32%). Removing SNPs and indels that were only variable compared to H37Rv and did not vary within the outbreak resulted in 278 SNPs and 92 indels, which was notable for the higher proportion of indels that were retained. Of these, 61 (21.9%) SNPs and 47 (51.1%) indels were phylogenetically informative and contributed to evolutionary signal. Indel polymorphisms were predominately found within gene regions (83 genic indels, 9 intergenic indels) and were evenly distributed along the genome (**Supplementary figure S1**). The numbers of insertions and deletions were similar (43 insertions and 49 deletions), and 37 of 92 indels were singletons (only found in one isolate in the outbreak). Annotation showed that most indels found outside of the PE/PPE gene family were in unannotated genes or hypothetical proteins (**Supplementary table S1**), with the most variation found in the *plcB* gene, which has previously been associated with possible changes in pathogenicity (Vera-Cabrera et al. 2007). There were no polymorphic indels in loci that have been previously associated with drug resistance.

Maximum likelihood phylogenetic reconstruction with SNP data showed a positive correlation between root-to-tip distance and collection date, with moderate temporal signal (R^2^ = 0.17, correlation coefficient = 0.41) (**Supplementary figure S2**). The temporal signal from the phylogenetic tree built with indels alone was weak, though with some positive correlation (R^2^ = 2.56 × 10^−2^, correlation coefficient 0.16). The phylogenetic tree built with SNP and indel data in partition was also found to have a weak positive correlation between divergence and collection date (R^2^ = 4.29 × 10^−2^, correlation coefficient 0.21). From the BEAST2 analysis to reconstruct timed phylogenies, the mutation rate of SNPs and indels across the outbreak was relatively even (**Supplementary table S2**), with 0.21 SNPs per year (HPD 0.17 – 0.25 SNPs) and 0.38 indels per year (HPD 0.31 – 0.44). The SNP mutation rate was estimated to be slightly higher in the SNP only analysis, with the estimated rate of 0.31 SNPs per year (HPD 0.20 – 0.44 SNPs).

### SNP and indel diversity of outbreak strains

This complex London outbreak of INH-resistant *Mtb* isolates is characterised by low SNP diversity and many isolates with identical sequences (Casali et al. 2016; Xu et al. 2020). There were 197 isolates with no SNP differences with at least one other sequence in the outbreak, with the largest single cluster consisting of 133 isolates with the same SNP profile (**Figure 1A**). This clonal cluster is marginally larger than identified in previous studies of this outbreak (Casali et al. 2016; Xu et al. 2020). Including indel polymorphism increased the detectable diversity between strains in the outbreak; the number of isolates with an identical sequence to at least one other isolate in the outbreak decreased to 113 isolates. The large clonal SNP cluster was broken up into four smaller clusters, the largest of which consisting of 63 sequences, along with 34 unique sequences (**Figure 1B**).

**Figure 1.**
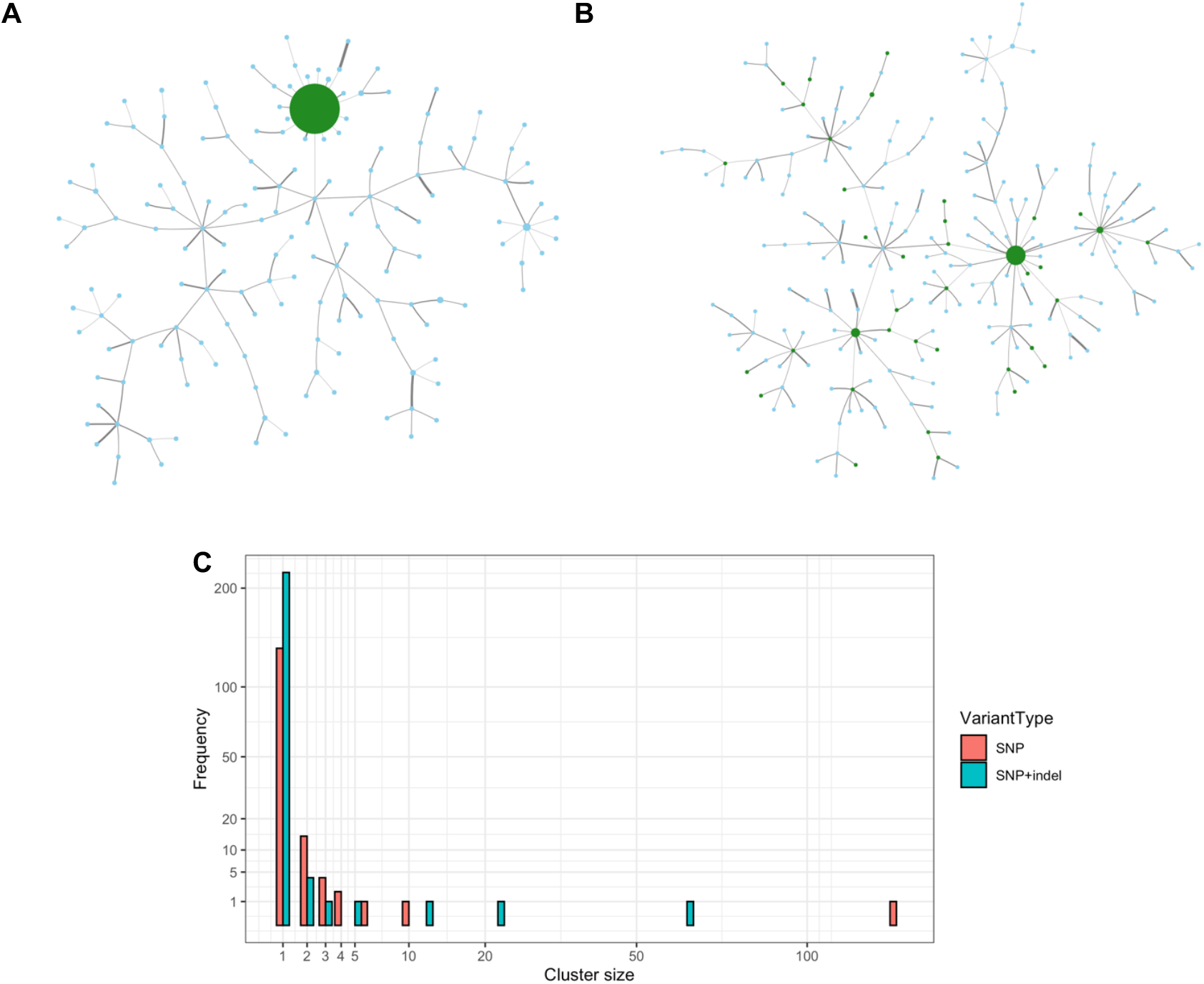
Network plot of clusters of identical sequences in the outbreak with node size relative to the cluster size and edge thickness as the SNP distance. (A) is constructed using SNP variation with the largest clonal SNP cluster (n = 133) is coloured green and all other clusters blue. (B) is constructed using the joint SNP and indel variation. The sequences in the largest clonal SNP cluster in this analysis are coloured green, with all other sequences coloured blue. (C) the cluster size distribution of identical sequences by SNPs and SNPs and indels. Axes have been scaled by the square root.

The maximum pairwise SNP distance between outbreak strains was 16 SNPs, with most isolates separated few SNPs (**Figure 2A**). The maximum pairwise indel distance between isolates was also 16, though the median pairwise indel distance was lower than the median SNP distance (3 SNPs (IQR 2 – 5 SNPs) vs 2 indels (IQR 1 – 3 indels)). There was a positive relationship between pairwise indel and SNP distance (Pearson’s correlation coefficient t = 16.84, P < 0.05) (**Figure 2B**), yet there were multiple cases of pairs with a low SNP distance and high indel distance. An extreme example was two strains in the outbreak that differed by only 1 SNP but had a pairwise indel distance of 16. Closer inspection of the sequence assemblies in these strains did not reveal an excess of missing or heterogeneous calls that may erroneously decrease distances.

**Figure 2.**
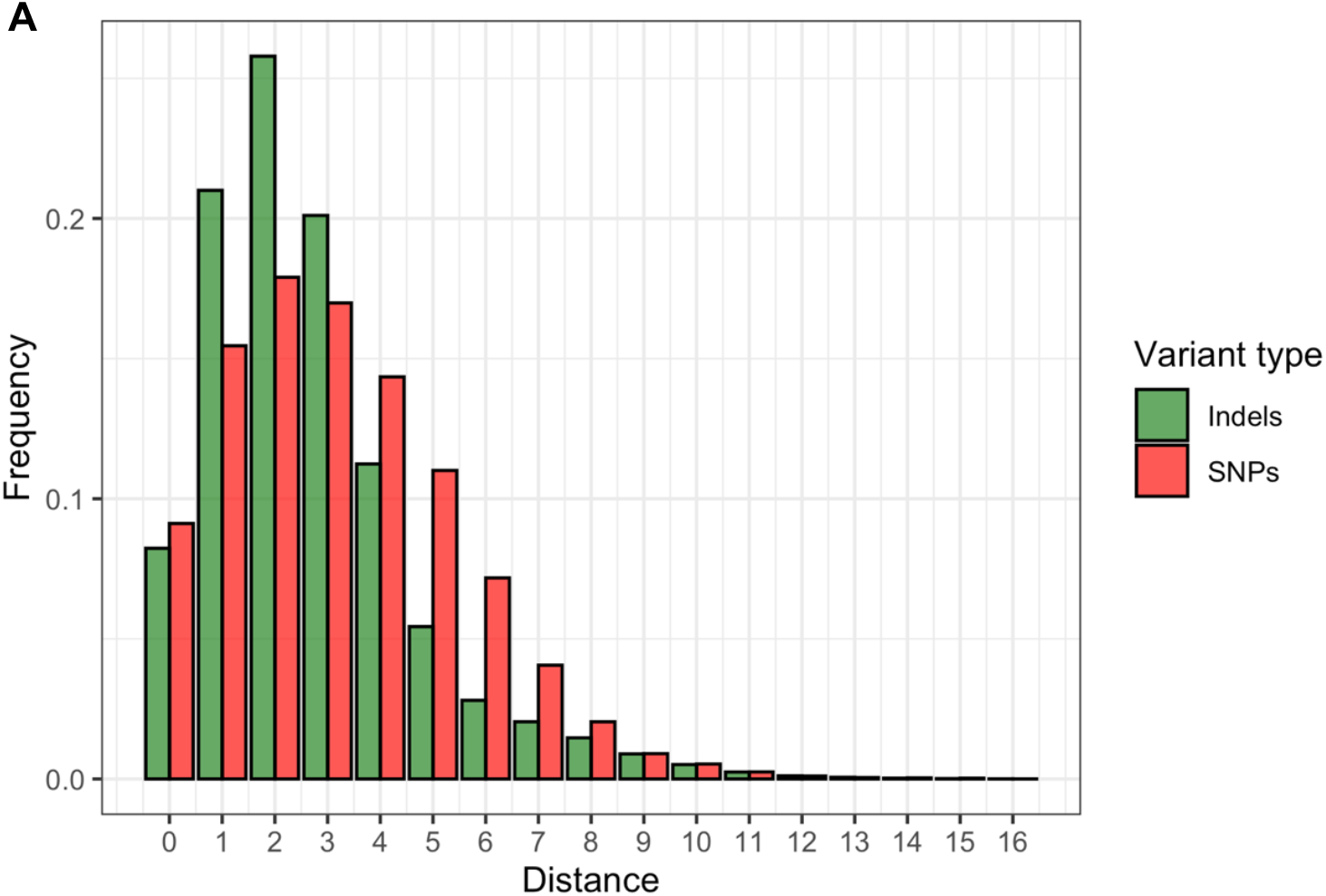
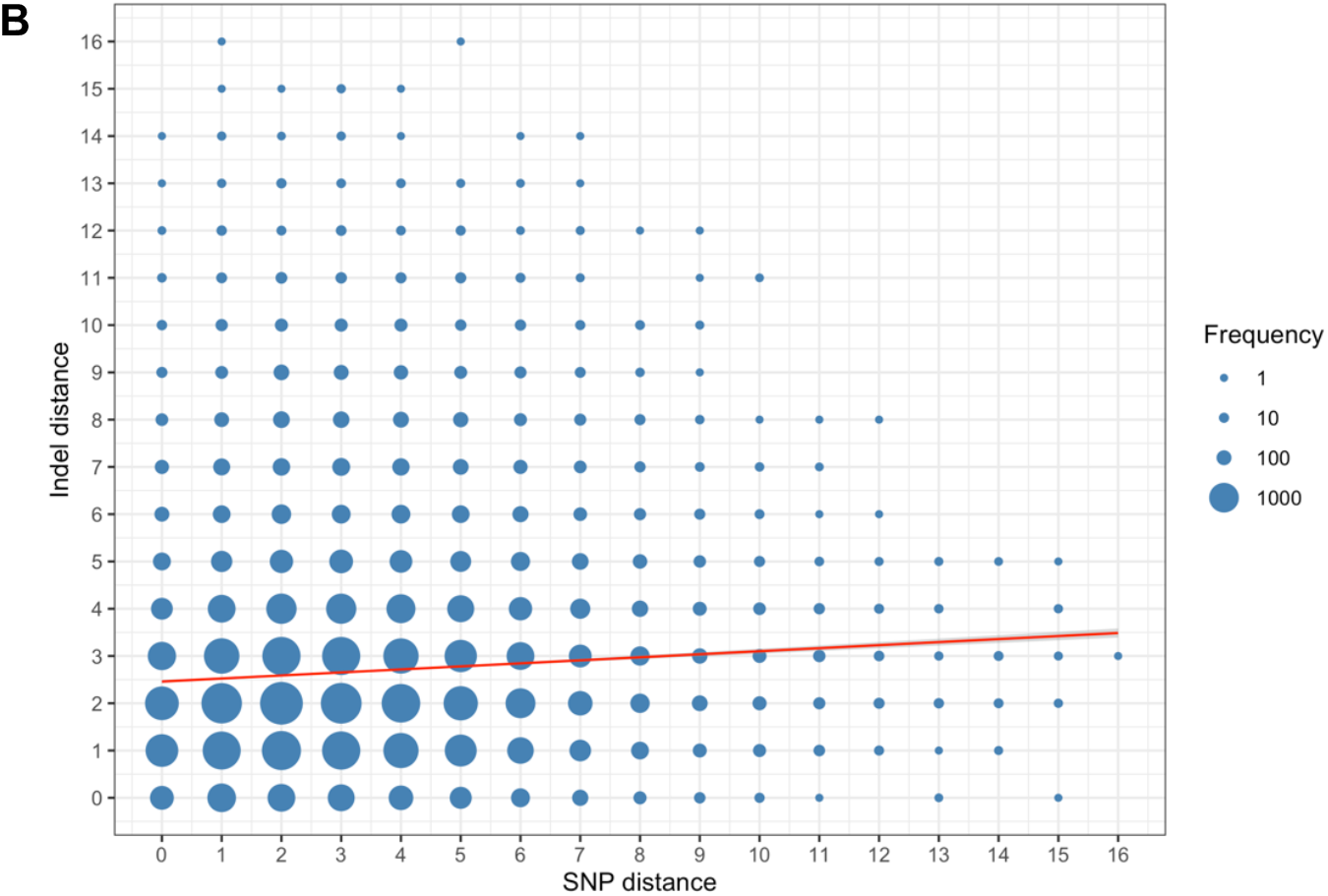
**(A)** A histogram of the frequency of pairwise SNP distances and pairwise indel distances, and **(B)** a scatter plot comparing pairwise SNP and indel distances between all *M. tuberculosis* isolates in the outbreak. The frequency of the pairwise distances between isolates is denoted by the size of each datapoint. A significant positive relationship (*P* < 0.05) is shown by linear regression.

### Transmission reconstruction including indel polymorphism

Transmission clusters defined using a pairwise SNP threshold of ≤ 5 SNPs resulted in a single large cluster with all but one isolate, reflecting the very low diversity within the outbreak (**Supplementary table S3**). There was little further delineation of this large cluster when including a threshold on the pairwise indel distance between isolates, with all sequences remaining in the cluster at thresholds of ≤ 5 and ≤ 3 indels, and all but three strains in the large cluster at the ≤ 1 indel threshold.

Direct transmission events between sampled hosts in the outbreak were inferred using TransPhylo (Didelot et al. 2017) from the timed phylogenies produced from SNPs, and from SNPs and indels jointly (**Figure 3**). The highest posterior probability between two samples was used to infer the direction of transmission. The resulting number of transmission events inferred from SNP data alone and from SNPs and indels varied depending on the probability threshold used to accept a transmission link, though the predicted number of events was similar across all probability thresholds between 0.1 and 1. (**Figure 3A)**.

**Figure 3.**
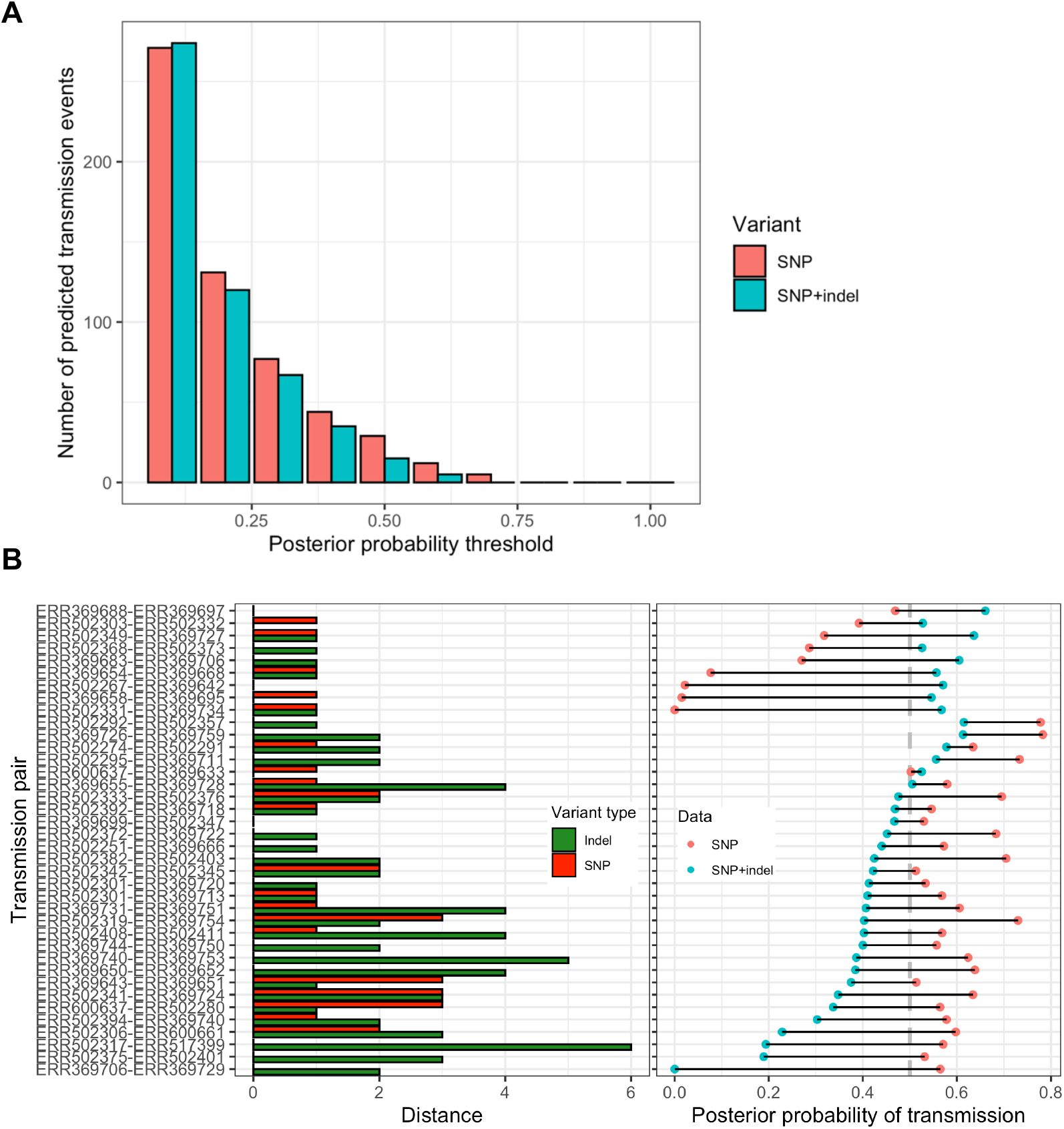
**A)** The number of inferred transmission events predicted using TransPhylo from SNP data (red) and SNP and indel data (blue) at a given probability threshold between 0.1 and 1, in increments of 0.1. Links with a posterior probability of < 0.1 are not shown. **B)** The right panel shows the likelihood of transmission events inferred using SNP data against the probability of the transmission event inferred using SNP and indel data together. Only transmission links with a posterior probability of ≥ 0.5 in either analysis are shown. The left panel shows the pairwise SNP and pairwise indel divergence between strains in the inferred transmission event.

Using a probability threshold of ≥ 0.5 (a given link was present in greater than 50% of posterior transmission trees), which has been used previously as a threshold to accept transmission events in this approach (Sobkowiak et al. 2020; Xu et al. 2020), we found that the SNP only analysis identified 29 putative transmission events, eight more links than in previous analysis by Xu *et. al*. (Xu et al. 2020). Of these, only 6/29 were also predicted with a probability of ≥ 0.5 from the transmission inference using SNPs and indels (**Figure 3B**). A further 13 of the 29 remaining transmission events identified using SNPs were predicted with a similar probability in the SNP and indel analysis (posterior probability ≥ 0.4). However, there were links with a significantly higher probability in the SNP analysis than with SNPs and indels. These transmission pairs typically had a low pairwise SNP distance and higher indel distance, though there were also transmission pairs with a low SNP distance and higher indel distance where the probability of direct transmission was predicted to be similar in the two analyses. Finally, nine additional transmission events were predicted in the joint SNP and indel analysis that did not meet the probability threshold in the SNP only analysis. As expected, these pairs were characterised by a low pairwise SNP and indel distance, increasing the probability of direct transmission in the joint analysis as these pairs remained close in the input phylogeny while sequences with a higher number of indel differences were more distant.

### Within-host indel evolution

Finally, to gain insights into the within-host indel diversity in the TB outbreak, we analysed sequence data from 12 individuals that were sampled twice over a single episode of TB disease. Interestingly, we found relatively high levels of both SNP and indel diversity between these samples taken from the same host, with a maximum of 4 SNP and 8 indel differences within any host that was sampled multiple times (**Figure 4A**). We did not find any significant association between SNP distance and indel distance in these serially sampled individuals (R^2^ = 0.01, *P* > 0.05). Additionally, there was no association between either SNP distance (R^2^ = 0.01, *P >* 0.05) or indel distance (R^2^ = 0.01, *P* > 0.05) and the difference between collection dates, suggesting that the diversity identified in these individuals is not driven by clock-like evolution within the host (**Figure 4B**).

**Figure 4.**
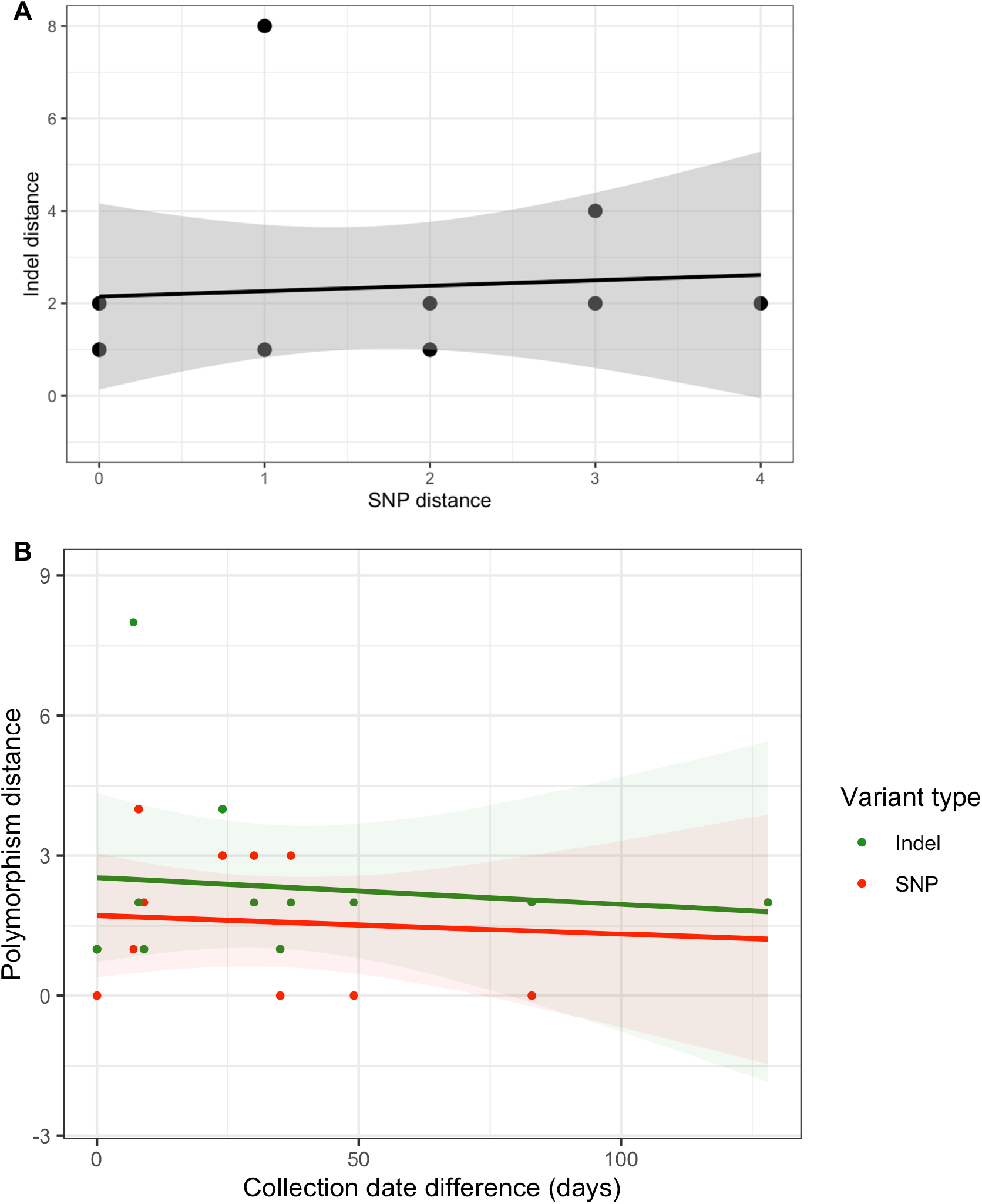
(A) Indel distance against SNP distance, and (B) Indel distance and SNP distance against the difference in collection date in 12 individuals sampled twice in a same TB disease episode, including the regression line of best fit and standard error. Some individual points will be overlain where distances are equal.

## Discussion

We present an analysis of the variation in short insertions and deletions in *Mtb* strains from a large, low diversity TB outbreak and have demonstrated how indel polymorphisms can be included to reconstruct transmission events from WGS data. We found that including indel variation reduced the number of identical sequences in this outbreak and altered the posterior probability of inferred transmission events compared to using SNP data alone. This finding can help to refine *Mtb* transmission networks reconstructed from WGS data and improve our understanding of the transmission dynamics in TB outbreaks.

Including indel variation between strains when predicting clusters of recent transmission in the outbreak did not markedly increase the resolution of clusters identified with the routinely used threshold of ≤ 5 SNPs, though indel variation does break up clusters of isolates that are indistinguishable using SNPs alone. The three tested thresholds for the indel distance, including a very stringent distance of up to one indel difference between sequences, only removed a small number of isolates from the large cluster of almost all sequences identified with the 5 SNP threshold. This suggests that within very closely related outbreaks such as we’ve investigated in this study, the variation added by indel polymorphisms may not be sufficient to delineate closer sub-clusters of recent transmission. Even so, we can increase the observed sequence diversity in an outbreak by including genomic variation from indels and reduce the number of identical sequences compared to sequence analysis using only SNP differences. In this analysis, the largest cluster of sequences that appear identical by SNPs was 133 isolates. Attempting to infer who-infected-whom in this cluster from genomic data and collection time, with the earliest strain as the source of infection, we would have 8778 equally probable unique pairs. In contrast, the largest cluster of identical sequences when including indel variation decreased to 63 isolates, which reduces the number of equally probable unique pairs by almost 80% to 1953 pairs. This can have great implications for phylogenetic reconstruction and the prediction of person-to-person transmission events due to an incorrect inference on the divergence between pairs if we do not include the substantial variation provided by indels.

This is evident in the differences in the transmission histories of the London outbreak from analysis using SNPs alone and when including indels. Examining direct transmission events inferred using TransPhylo from phylogenies produced with SNP data and with SNPs and indels jointly, we found a comparative number of predicted links between pairs. Nonetheless, there were key differences found in high-confidence transmission events (posterior probability ≥ 0.5) between the two analyses that can be attributed to identifying indel polymorphisms in the transmission pair. All but one of the transmission events only predicted using SNP data that did not reach the probability threshold when including indels in the analysis were between strains with at least one indel polymorphism. The three links with the greatest difference between posterior probability of direct transmission estimated from SNPs and from SNPs and indels (0.34 – 0.56 decrease in probability) were between strains with no SNP variation but with a pairwise distance of between two and six indels. There were also relatively large differences between the posterior probability of transmission events between isolates that were often characterised by pairs with a low SNP distance and a higher indel distance.

However, there were also cases where there were differences between the predicted probability of direct transmission in the two analyses with low pairwise indel distance between strains. This included transmission events predicted with a high probability in the SNP and indel joint analysis that were predicted with a low probability in the transmission reconstruction using only SNP variation. The posterior probability in TransPhylo is calculated as the proportion of the posterior transmission trees that contained a given transmission event, and so these values may differ between analyses for two reasons. Firstly, the posterior distribution of inferred transmission trees will change with different MCMC runs, and even if the model reaches convergence, there may be minor differences in the posterior probability of transmission chains. For this reason, minor differences in the posterior probabilities of transmission between a pair may mean it is not accepted if it falls just below the given threshold. Instead, it may be advantageous to use the probability value itself for any downstream analysis of transmission dynamics rather than a simple threshold. Secondly, increasing the divergence between a pair by including indel variation will reduce the likelihood of these strains being placed close to each other in a phylogeny. Accordingly, this will increase the predicted probability of direct transmission with other strains closer on the tree where there are no indel polymorphisms between the pair.

The temporal evolutionary signal found in indels within the outbreak strains was weaker than from SNPs alone, as determined during phylogenetic tree building with distinct substitution models. However, there was still a weak positive correlation between time and genetic distance when genetic distance was measured with SNPs and indels jointly. The added variation that is detected when indels are included can reveal evolutionary differences between strains that may appear identical or near identical when considering only SNP diversity, as shown in our data. A clock-like evolution of indels has been shown previously with the analysis of a multidrug resistant *Mtb* outbreak (Godfroid et al. 2020), with an enrichment of indels in known drug resistance associated genes. Interestingly in our dataset, we identified only one drug resistance loci in indels in the London isolates when compared to the H37Rv reference strain. This was a deletion in the *rrs* gene that has been previously associated with streptomycin resistance (Tudó et al. 2010), though this was present within all strains in the outbreak. This suggests that potential neutral or near neutral indel evolution across the genome in the absence of strong selection increases the genomic diversity between strains at this resolution.

We also examined the within host diversity over time by characterising the genomic variation, both SNPs and indels, in 12 individuals that were sampled at two timepoints across the same disease episode. We found no significant association between the number of SNP or indel polymorphisms in samples from the same host over time, and no association between SNP distance and indel distance. As such, we found no evidence of within host sequence evolution through time in these strains. Instead, these results likely reflect inherent issues with sample preparation and sequencing from culture, which can introduce genetic bottlenecks through differential growth in culture media and the mechanism of sampling microbial communities that may not capture the full genetic diversity (Shockey et al. 2019). Sequencing *Mtb* directly from sputum, which has shown to increase the detectable diversity in isolates (Nimmo et al. 2019), would allow for a better representation of the complete within host *Mtb* genomic diversity in a sample. Additionally, the presence of at least one indel polymorphism in multiple samples from the same host, even at the same timepoint, may also indicate there are issues with the accuracy of indel variant calling from the short read sequence data used in this study. Long read sequencing can improve detection in difficult to assemble regions, such as indels (De Coster et al. 2021), which will improve the analysis of indel diversity and evolution in *Mtb* outbreaks.

There were limitations to the analysis performed in this study. The transmission reconstruction methodology involves producing a timed phylogeny before inferring transmission trees, which is a hypothesis about the phylogenetic relationships between isolates. To account for this phylogenetic uncertainty, we used the TransPhylo approach that takes multiple input phylogenies and jointly estimates transmission trees and ensured convergence across all posterior parameters in the tree building process. We also performed model testing prior to building the timed phylogenies to identify the optimal substitution models for the data. Even so, the models available were not optimised for indels, which are currently not well established due to complexity (Loewenthal et al. 2021), and advances in developing these models and implementation in tree building software will improve phylogenetic tree inference including these variants. A further limitation was the absence of a ‘gold-standard’ validation of the predicted transmission events to confirm true links and evaluate the accuracy of the transmission reconstruction that incorporated indel polymorphisms against SNP only data. Obtaining these confirmed links is difficult as comprehensive contact investigation data are required, which are often not available for all pairs in an outbreak and can often disagree with genomic inferences (Alaridah et al. 2019; Guthrie et al. 2020). Nevertheless, our finding that the greatest change in the posterior probability of transmission events between the two analyses was in pairs with indel polymorphisms indicates that including these variants can increase the informative sequence diversity used to infer true transmission.

Here, we have shown that variation in short insertions and deletions can be included in transmission reconstruction from WGS data to provide additional diversity between isolates. Including variation from indels can refine transmission reconstruction by increasing the observed divergence between strains to differentiate sequences that appear identical by SNPs. This can also change the inferred transmission network using probabilistic models where the relative genomic distance between isolates in a population is adjusted. Further work on sequence assembly and variant calling of indels and advances in developing optimal indel substitution rates for phylogenetic reconstruction would improve this analysis, increasing the utility of indel polymorphisms in transmission analysis from sequencing data.

## Methods and Materials

### Sample data and sequence analysis

Samples were collected as part of well-characterised INH-resistant TB outbreak in London, UK, designated cluster E1244 by Public Health England. All isolates were collected between 2006 – 2013 and details on the data collection and previous epidemiological investigations of the cluster are described elsewhere (Ruddy et al. 2004; Neely et al. 2010; Maguire et al. 2011; Casali et al. 2016). Raw WGS data are available from the European Nucleotide Archive (accession number ERP003508), and culturing and sequencing information is detailed in Xu *et. al*. (Xu et al. 2020).

Sequences were inspected for quality using FastQC (v.0.11.9) and trimmed and filtered using Trimmomatic (Bolger et al. 2014). A reference-guided alignment was performed using BWA ‘mem’ (Li and Durbin 2009) to map reads against the H37Rv reference strain (NC_000962.3) and resulting alignment files sorted using SAMtools (Li et al. 2009). SNPs and indels were called using the GATK ‘HaplotypeCaller’ and ‘GenetypeGVCFs’ tools. Recent developments in this software have improved the detection of short indels from short read sequence data (Wang et al. 2022) and all final indel polymorphisms identified were validated using Pilon (Walker et al. 2014). Poor quality and low coverage variants (Phred score < 20 and read depth < 5) were removed, and SNPs and indels in repeat regions were excluded. Heterozygous calls with a support of > 80% of reads were called as the consensus allele, otherwise the site was called as the ambiguous call ‘N’. All samples were tested using MixInfect (Sobkowiak et al. 2018) and found to contain a single strain.

### Phylogenetic analysis

We used ModelFinder (Kalyaanamoorthy et al. 2017), part of the IQ-TREE package (v.1.6.12) (Minh et al. 2020), to determine the best-fit substitution and morphological models for phylogenetic reconstruction through the best scoring Akaike information criterion (AIC) and Bayesian information criterion (BIC). ModelFinder found the substitution model ‘TVM’ with equal base frequencies and ascertainment correction bias (‘TVMe+ASC’) to be the optimal substitution model in our SNP data and the Jules-Cantor type model with equal frequencies, ascertainment bias, and a FreeRate model with four categories (‘ MK+FQ+ASC+R4’) as the optimal model for the indel data. Indels were treated as a morphological characteristic rather than binary presence/absence, as multiple length insertions and deletions can be present between samples at the same locus. All sequences passed the composition chi-square test to check for homogeneity of nucleotide and site composition. The resulting maximum-likelihood phylogenies produced by IQ-TREE were used to explore the temporal signal in the SNP and indel data using TempEst (v.1.5.3) (Rambaut et al. 2016). We also performed this analysis with a partitioned SNP and indel dataset to investigate the joint signal when considering both forms of genomic variation together.

Time-calibrated phylogenetic trees were produced using BEAST2 (v.2.6.6) (Bouckaert et al. 2019) for SNP data and SNP and indels together as partitioned input data with a linked tree. Optimal substitution models identified with ModelFinder were applied in BEAST2 for SNP and indel data by installing the standard substitution models (SSM) package for SNP data (Bouckaert and Xie 2017 Sep 25), and the morphological models (MM) package for indel data (Lewis 2001). As there was a temporal signal found in SNPs, we used a strict molecular clock with an initial value of 1×10^−7^ substitutions per site per year (Ford et al. 2013), while a lognormal relaxed molecular clock was used for indels, as the temporal signal was weaker in these data. We used a constant population size with a lognormal prior distribution and corrected for invariant sites in SNP data by manually adapting the XML files, as described previously (Xu et al. 2020). BEAST2 was run for 5×10^8^ MCMC iterations, sampling every 1000^th^ tree until convergence, which was heuristically indicated when all parameters reached an effective sample size (ESS) of ≥ 200 in Tracer (Rambaut et al. 2018). A maximum clade credibility tree for each run was produced from the posterior set of trees using TreeAnnotator in BEAST2, with a 20% burn-in, and visualised using iTOL (Letunic and Bork 2021).

### Transmission analysis

Putative genomic transmission clusters were identified from pairwise SNP and indel distances by linking isolates that were within a given threshold. As has been used previously to define recent transmission (Meehan et al. 2019; Nikolayevskyy et al. 2019; Cancino-Muñoz et al. 2022), we employed a threshold of 5 SNPs cluster isolates and tested the threshold for indel distance at 1, 3, and 5 indels. We then used TransPhylo (Didelot et al. 2017) to reconstruct the transmission network in the London *Mtb* outbreak. TransPhylo employs a Bayesian framework to infer the probability of transmission events from phylogenetic trees. Specifically, we used the recent extension of TransPhylo that accounts for phylogenetic uncertainty by simultaneously inferring transmission from a sample of the posterior timed trees produced by BEAST2, rather than a single consensus tree, with model parameters shared across trees resulting in single inference from the multiple input trees (Xu et al. 2020). We chose a random sample of 50 posterior input trees after a 50% burn-in, running TransPhylo for 1.0*e*^6^ iterations. Epidemiological parameters, including the generation time distribution and sampling density, were chosen as per Xu *et. al*. (Xu et al. 2020). We conducted two transmission analyses, first using the phylogenies produced using SNP data alone and then with the phylogenies produced from the combined SNP and indel data in partition, inspecting differences between the inferred transmission events.

## Acknowledgments and funding information

We would like to thank Vijay Naidu for compiling the sequence data and useful discussions. This work was supported by the Federal Government of Canada’s Canada 150 Research Chair program.

---

**Supplementary figure S1.**
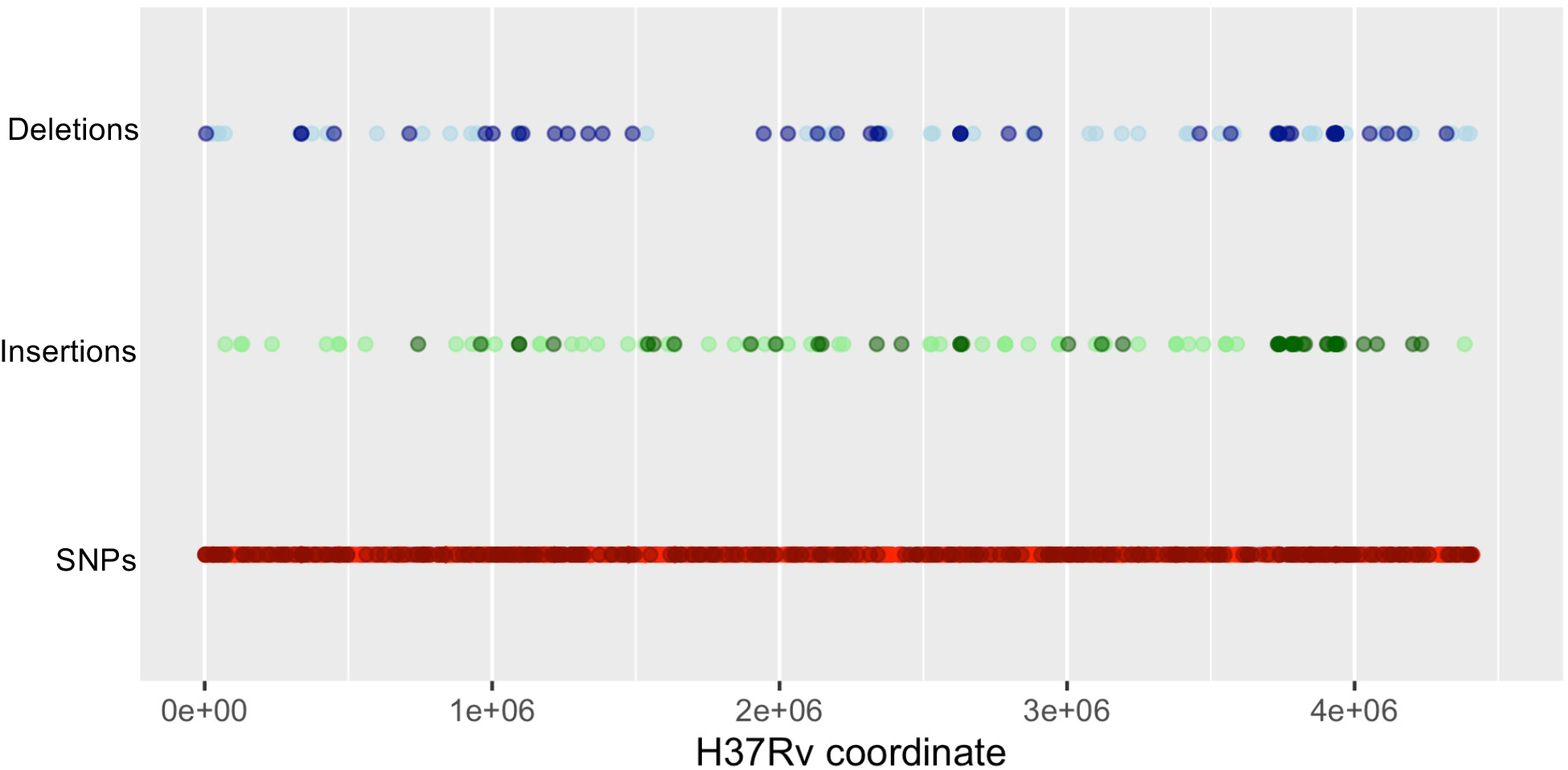
The positions of the SNP and indel variants identified in the London outbreak, indexed by the H37Rv reference strain (NC_000962.3). SNPs are shown in red, deletions in blue and insertions in green. Darker colours denote polymorphism that vary within outbreak strains and lighter colours are those that vary only in respect to H37Rv but are invariant within the outbreak strains.

**Supplementary figure S2.**
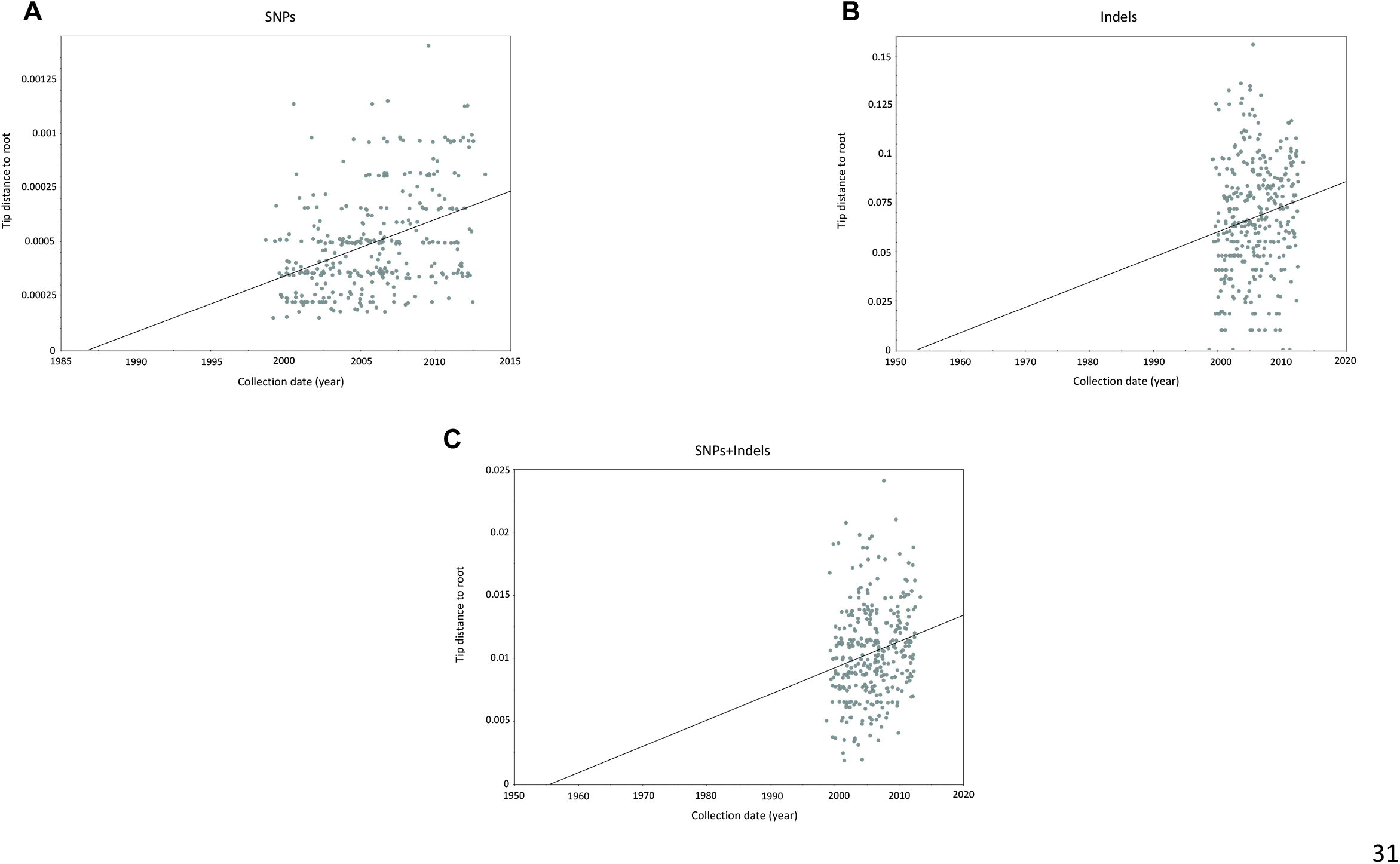
Scatter plots showing the correlation between root-to-tip distances in the maximum-likelihood phylogeny and collection dates of the London *M. tuberculosis* outbreak strains. Trees were constructed from A) SNPs only, B) indels only, and C) SNPs and indels jointly. Plots were produced using TempEst and each datapoint is a single sample, with the best-fit regression line drawn on each plot.

**Supplementary Table S1 (Excel spreadsheet).** Annotation of the indel polymorphisms called in 332 *Mycobacterium tuberculosis* isolates from London, UK. Variation in PE/PPE genes, repetitive regions, and known antimicrobial resistance genes were removed, along with indels that were invariant between the samples in the outbreak.

**Supplementary Table S2.** The posterior clock rate and tree height estimates from the BEAST phylogenetic analysis using different data sources to build phylogenies: SNPs only, and SNPs and indels jointly.

| Input data | Mean clock rate –<br>substitutions per site<br>per year<br>(95% HPD) | Polymorphisms per<br>year<br>(95% HPD) | Tree height in years<br>(95% HPD) |
| --- | --- | --- | --- |
| SNPs | $7.0 \times 10^{-8}$<br>($4.5 \times 10^{-8} - 1.0 \times 10^{-7}$)* | 0.31 SNPs<br>(0.20 – 0.44 SNPs) | 27.7 (21.6 – 34.7) |
| SNPs &<br>indels (SNPs) | $4.7 \times 10^{-8}$<br>($3.9 \times 10^{-8} - 5.6 \times 10^{-8}$)* | 0.21 SNPs<br>(0.17 – 0.25 SNPs) | 34.0 (26.6 – 42.3) |
| SNPs &<br>indels (indels) | $4.1 \times 10^{-3}$<br>($3.4 \times 10^{-3} - 4.8 \times 10^{-3}$) | 0.38 indels<br>(0.31 – 0.44 indels) | |

\*Corrected for invariant sites

**Supplementary table S3 (Excel Spreadsheet):** Genomic cluster designation of the 332 *Mycobacterium tuberculosis* isolates, based on 5 a pairwise threshold of 5 SNPs and any number of indels, 5 SNPs and 5 indels, 5 SNPs and 3 indels, and 5 SNPs and 1 indel.

## References

Alaridah N, Hallbäck ET, Tångrot J, Winqvist N, Sturegård E, Florén-Johansson K, Jönsson B, Tenland E, Welinder-Olsson C, Medstrand P, et al. 2019. Transmission dynamics study of tuberculosis isolates with whole genome sequencing in southern Sweden. Sci Rep. 9(1):4931.

Ayabina D, Ronning JO, Alfsnes K, Debech N, Brynildsrud OB, Arnesen T, Norheim G, Mengshoel A-T, Rykkvin R, Dahle UR, et al. 2018. Genome-based transmission modelling separates imported tuberculosis from recent transmission within an immigrant population. Microb Genomics. 4(10):1–13.

Bolger AM, Lohse M, Usadel B. 2014. Trimmomatic: A flexible trimmer for Illumina sequence data. Bioinformatics. 30(15):2114–2120.

Bouckaert R, Vaughan TG, Barido-Sottani J, Duchêne S, Fourment M, Gavryushkina A, Heled J, Jones G, Kühnert D, De Maio N, et al. 2019. BEAST 2.5: An advanced software platform for Bayesian evolutionary analysis. PLoS Comput Biol. 15(4):1–28.

Bouckaert R, Xie D. 2017 Sep 25. BEAST2-Dev/substmodels: Standard Nucleotide Substitution Models v1.0.1. Zenodo. doi:10.5281/ZENODO.995740.

Cancino-Muñoz I, López MG, Torres-Puente M, Villamayor LM, Borrás R, Borrás-Máñez M, Bosque M, Camarena JJ, Colijn C, Colomer-Roig E, et al. 2022. Population-based sequencing of Mycobacterium tuberculosis reveals how current population dynamics are shaped by past epidemics. Elife. 11:1–23.

Casali N, Broda A, Harris SR, Parkhill J, Brown T, Drobniewski F. 2016. Whole Genome Sequence Analysis of a Large Isoniazid-Resistant Tuberculosis Outbreak in London: A Retrospective Observational Study. PLoS Med. 13(10):1–18.

Coll F, Preston M, Guerra-Assunção JA, Hill-Cawthorn G, Harris D, Perdigão J, Viveiros M, Portugal I, Drobniewski F, Gagneux S, et al. 2014. PolyTB: A genomic variation map for Mycobacterium tuberculosis. Tuberculosis. 94(3):346–354.

De Coster W, Weissensteiner MH, Sedlazeck FJ. 2021. Towards population-scale long-read sequencing. Nat Rev Genet. 22(9):572–587.

Didelot X, Fraser C, Gardy J, Colijn C. 2017. Genomic infectious disease epidemiology in partially sampled and ongoing outbreaks. Mol Biol Evol. 34(4):997–1007.

Ford CB, Shah RR, Maeda MK, Gagneux S, Murray MB, Cohen T, Johnston JC, Gardy J, Lipsitch M, Fortune SM. 2013. Mycobacterium tuberculosis mutation rate estimates from different lineages predict substantial differences in the emergence of drug-resistant tuberculosis. Nat Genet. 45(7):784–790.

Gardy JL, Johnston JC, Sui SJH, Cook VJ, Shah L, Brodkin E, Rempel S, Moore R, Zhao Y, Holt R, et al. 2011. Whole-Genome Sequencing and Social-Network Analysis of a Tuberculosis Outbreak. N Engl J Med. 364(8):730–739.

Godfroid M, Dagan T, Merker M, Kohl TA, Diel R, Maurer FP, Niemann S, Kupczok A. 2020. Insertion and deletion evolution reflects antibiotics selection pressure in a mycobacterium tuberculosis outbreak. PLoS Pathog. 16(9):1–24.

Gupta A, Alland D. 2021. Reversible gene silencing through frameshift indels and frameshift scars provide adaptive plasticity for Mycobacterium tuberculosis. Nat Commun. 12(1):1–11.

Guthrie JL, Strudwick L, Roberts B, Allen M, McFadzen J, Roth D, Jorgensen D, Rodrigues M, Tang P, Hanley B, et al. 2020. Comparison of routine field epidemiology and whole genome sequencing to identify tuberculosis transmission in a remote setting. Epidemiol Infect. 148:e15.

Hatherell HA, Didelot X, Pollock SL, Tang P, Crisan A, Johnston JC, Colijn C, Gardy JL. 2016. Declaring a tuberculosis outbreak over with genomic epidemiology. Microb Genomics. 2(5):3–6.

Kalyaanamoorthy S, Minh BQ, Wong TKF, Von Haeseler A, Jermiin LS. 2017. ModelFinder: Fast model selection for accurate phylogenetic estimates. Nat Methods. 14(6):587–589.

Klinkenberg D, Backer JA, Didelot X, Colijn C, Wallinga J. 2017. Simultaneous inference of phylogenetic and transmission trees in infectious disease outbreaks.

Lee RS, Proulx JF, McIntosh F, Behr MA, Hanage WP. 2020. Previously undetected super-spreading of mycobacterium tuberculosis revealed by deep sequencing. Elife. 9:1–15.

Letunic I, Bork P. 2021. Interactive tree of life (iTOL) v5: An online tool for phylogenetic tree display and annotation. Nucleic Acids Res. 49(W1):W293–W296.

Lewis PO. 2001. A likelihood approach to estimating phylogeny from discrete morphological character data. Syst Biol. 50(6):913–925.

Li H, Durbin R. 2009. Fast and accurate short read alignment with Burrows-Wheeler transform. Bioinformatics. 25(14):1754–60.

Li H, Handsaker B, Wysoker A, Fennell T, Ruan J, Homer N, Marth G, Abecasis G, Durbin R. 2009. The Sequence Alignment/Map format and SAMtools. Bioinformatics. 25(16):2078–2079.

Loewenthal G, Rapoport D, Avram O, Moshe A, Wygoda E, Itzkovitch A, Israeli O, Azouri D, Cartwright RA, Mayrose I, et al. 2021. A Probabilistic Model for Indel Evolution: Differentiating Insertions from Deletions. Mol Biol Evol. 38(12):5769–5781.

Maguire H, Brailsford S, Carless J, Yates M, Altass L, Yates S, Anaraki S, Charlett A, Lozewicz S, Lipman M, et al. 2011. Large outbreak of isoniazid-monoresistant tuberculosis in London, 1995 to 2006:Case-control study and recommendations. Eurosurveillance. 16(13).

Meehan CJ, Goig GA, Kohl TA, Verboven L, Dippenaar A, Ezewudo M, Farhat MR, Guthrie JL, Laukens K, Miotto P, et al. 2019. Whole genome sequencing of Mycobacterium tuberculosis: current standards and open issues. Nat Rev Microbiol. 17(9):533–545.

Minh BQ, Schmidt HA, Chernomor O, Schrempf D, Woodhams MD, Von Haeseler A, Lanfear R, Teeling E. 2020. IQ-TREE 2: New Models and Efficient Methods for Phylogenetic Inference in the Genomic Era. Mol Biol Evol. 37(5):1530–1534.

Neely F, Maguire H, Le Brun F, Davies A, Gelb D, Yates S. 2010. High rate of transmission among contacts in large London outbreak of isoniazid mono-resistant tuberculosis. J Public Health (Bangkok). 32(1):44–51.

Nikolayevskyy V, Niemann S, Anthony R, van Soolingen D, Tagliani E, Ködmön C, van der Werf MJ, Cirillo DM. 2019. Role and value of whole genome sequencing in studying tuberculosis transmission. Clin Microbiol Infect. 25(11):1377–1382.

Nimmo C, Shaw LP, Doyle R, Williams R, Brien K, Burgess C, Breuer J, Balloux F, Pym AS. 2019. Whole genome sequencing Mycobacterium tuberculosis directly from sputum identifies more genetic diversity than sequencing from culture. BMC Genomics. 20(1):1–9.

Phelan JE, O’Sullivan DM, Machado D, Ramos J, Oppong YEA, Campino S, O’Grady J, McNerney R, Hibberd ML, Viveiros M, et al. 2019. Integrating informatics tools and portable sequencing technology for rapid detection of resistance to anti-tuberculous drugs. Genome Med. 11(1):1–7.

Rambaut A, Drummond AJ, Xie D, Baele G, Suchard MA. 2018. Posterior summarization in Bayesian phylogenetics using Tracer 1.7. Syst Biol. 67(5):901–904.

Rambaut A, Lam TT, Carvalho LM, Pybus OG. 2016. Exploring the temporal structure of heterochronous sequences using TempEst (formerly Path-O-Gen). Virus Evol. 2(1):1–7.

Romanowski K, Sobkowiak B, Guthrie JL, Cook VJ, Gardy JL, Johnston JC. 2020. Using Whole Genome Sequencing to Determine the Timing of Secondary Tuberculosis in British Columbia, Canada. Clin Infect Dis. 50(9):1052–1063.

Ruddy MC, Davies AP, Yates MD, Yates S, Balasegaram S, Drabu Y, Patel B, Lozewicz S, Sen S, Bahl M, et al. 2004. Outbreak of isoniazid resistant tuberculosis in north London. Thorax. 59(4):279–285.

Shockey AC, Dabney J, Pepperell CS, Tobin DM, Gordon S V. 2019. Effects of Host, Sample, and in vitro Culture on Genomic Diversity of Pathogenic Mycobacteria. 10(June):1–14.

Sobkowiak B, Banda L, Mzembe T, Crampin AC, Glynn JR, Clark TG. 2020. Bayesian reconstruction of Mycobacterium tuberculosis transmission networks in a high incidence area over two decades in Malawi reveals associated risk factors and genomic variants. Microb Genomics. 6(4).

Sobkowiak B, Glynn JR, Houben RMGJ, Mallard K, Phelan JE, Guerra-Assunção JA, Banda L, Mzembe T, Viveiros M, McNerney R, et al. 2018. Identifying mixed Mycobacterium tuberculosis infections from whole genome sequence data. BMC Genomics. 19(1):613.

Tudó G, Rey E, Borrell S, Alcaide F, Codina G, Coll P, Martín-Casabona N, Montemayor M, Moure R, Orcau À, et al. 2010. Characterization of mutations in streptomycin-resistant Mycobacterium tuberculosis clinical isolates in the area of Barcelona. J Antimicrob Chemother. 65(11):2341–2346.

Vera-Cabrera L, Molina-Torres CA, Hernández-Vera MA, Barrios-García HB, Blackwood K, Villareal-Treviño L, Ocampo-Candiani J, Welsh O, Castro-Garza J. 2007. Genetic characterization of Mycobacterium tuberculosis clinical isolates with deletions in the plcA–plcB–plcC locus. Tuberculosis. 87(1):21–29.

Walker BJ, Abeel T, Shea T, Priest M, Abouelliel A, Sakthikumar S, Cuomo CA, Zeng Q, Wortman J, Young SK, et al. 2014. Pilon: An integrated tool for comprehensive microbial variant detection and genome assembly improvement. PLoS One. 9(11).

Wang N, Lysenkov V, Orte K, Kairisto V, Aakko J, Khan S, Elo LL. 2022. Tool evaluation for the detection of variably sized indels from next generation whole genome and targeted sequencing data. PLoS Comput Biol. 18(2):1–27.

World Health Organization. 2017. The End Strategy TB. End TB Strateg. 53(9):1689–1699.

World Health Organization. 2021. Global tuberculosis report 2021. Licence: CC BY-NC-SA 3.0 IGO.

Xu Y, Stockdale JE, Naidu V, Hatherell H, Stimson J, Stagg HR, Abubakar I, Colijn C. 2020. Transmission analysis of a large tuberculosis outbreak in London: a mathematical modelling study using genomic data. Microb Genomics. 6(11).

